# Co-occurrence of multiple CRISPRs and *cas* clusters suggests epistatic interactions

**DOI:** 10.1101/592600

**Authors:** Aude Bernheim, David Bikard, Marie Touchon, Eduardo PC Rocha

**Author notes:** To whom correspondence should be addressed. Aude Bernheim.

## Abstract

Prokaryotes use CRISPR-Cas for adaptive immunity, but the reasons for the existence of multiple CRISPR and *cas* clusters remain poorly understood. We found that more than 40% of the genomes encoding a system show atypical genetic organizations. Their analysis revealed negative and positive epistatic interactions between Cas subtypes. The latter often result in one single complex locus with a shared adaptation module and diverse interference mechanisms, presumably to produce more effective immune systems. We typed CRISPRs that could not be unambiguously associated with a *cas* cluster and found that such complex loci tend to have unique type I repeats in multiple CRISPRs. In contrast, under-represented co-occurrences caused by functional interference or redundancy may lead to CRISPRs distant from *cas* genes. To investigate the origin of atypical CRISPR-Cas organizations, we analyzed plasmids and phages. Sets of nearly 2000 phages and 10000 prophages were almost devoid of CRISPR-Cas systems, but a sizeable fraction of plasmids had them. Isolated CRISPRs in plasmids were often compatible with the chromosomal *cas* clusters, suggesting that plasmids use CRISPRs to subvert host immunity. These results point to an important role for the interactions between multiple CRISPR and Cas in the function and evolution of bacterial immunity.

## Introduction

CRISPR-Cas are an adaptive immune system that protects bacterial and archaeal cells from exogenous mobile genetic elements, such as viruses^1–4^. They are composed of a CRISPR and a cluster of *cas* genes. CRISPRs comprise two types of sequences: repeats and spacers. Repeats are short sequences (typically 20-40 bp) identical within a CRISPR. They are interspaced by short and diverse spacer sequences (typically 20-40 bp), which often match sequences from mobile genetic elements. The number of repeats in a CRISPR can be an indicator of its activity, because an array with many spacers can target a larger number of mobile genetic elemets^5^. The cluster of *cas* genes encodes the proteins involved in the three stages of CRISPR-Cas immunity: expression, interference, and immunity^6^. During expression, the CRISPR is transcribed and then processed into smaller RNAs called crRNA (CRISPR RNA), each carrying sequences from a repeat and a single spacer. Each of these crRNA serves as a guide for a complex of Cas proteins. If the sequence of a guide is complementary to other DNA sequence in the cell, for example from a phage infecting the cell, the complex will activate an immune response. For most types of CRISPR-Cas systems this leads to the cleavage and degradation of the invading DNA. During adaptation, a complex of Cas proteins (including Cas1 and Cas2) generates and then incorporates a new spacer in the CRISPR^6,7^.

CRISPR-Cas systems are present in less than half of Bacteria and in most Archaea^8^. They are extremely diverse and have been classified hierarchically according to the composition of the cluster of *cas* genes. They are grouped in two classes, six types (I to VI) and more than 20 subtypes^8–10^. Novel types have been recently proposed ^11,12^, but they are rare and will thus not be analysed in this study. The last surveys of CRISPR-Cas systems abundance and diversity among fully sequenced bacterial genomes included 2740 and 2751 genomes^8,13^. Makarova and colleagues detected 1,949 distinct *cas* clusters and 4,210 CRISPRs from 1,302 genomes out of 2740. They could assign a subtype to 93% of the *cas* clusters. Similar results were found by other authors^13^. CRISPR-Cas systems are frequently horizontally transferred^8,14,15^. They have been detected on diverse mobile genetic elements (MGE) like plasmids, phages or transposons ^16–20^, but a quantitative analysis of the prevalence of these elements is lacking.

The ability of CRISPR-Cas systems to acquire new spacers makes them very versatile, and able to tackle a large diversity of MGEs. Nevertheless, some genomes have been found to encode several CRISPR-Cas systems^8^. This is intriguing: why should a genome encode more than one adaptive system? A functional interaction between these systems was recently demonstrated in *Marinomonas mediterranea*, which carries both a subtype I-F and a subtype III-B system. There, it was found that the type III-B system can use crRNAs from the type I-F system, enabling the same guide RNA to target phages with different interference modules. These different Cas interference complexes have diverse molecular requirements, thus limiting the emergence of phages escape mutants^21^. Many *cas* genes are found near CRISPRs, but distant arrays (i.e., CRISPRs without neighboring *cas* genes) have also been identified^8,22^. They can be processed by Cas proteins encoded in other regions of the genome (in *trans*)^7^ or they may represent remnant systems.

Even though CRISPR and Cas proteins are parts of one system and both elements are required for adaptation and interference, there have been few quantitative studies integrating information on both CRISPR and Cas proteins. Here, we analyse the joint distribution of CRISPR and *cas* genes in a large set of fully sequenced bacterial genomes and their mobile genetic elements. We focus on genomes and loci encoding several of these elements to understand why they co-occur. Our results reveal preferential associations between certain systems, sometimes in complex genetic loci that constitute one single CRISPR-Cas system with one adaptation and several types of interference modules.

## Material and methods

### Data

We analyzed 5775 complete genomes retrieved from NCBI RefSeq representing 2268 and 167 species of Bacteria and Archaea (http://ftp.ncbi.nih.gov/genomes/refseq/bacteria/), in November 2016. These genomes contained 4453 plasmids that were analysed in this work. Because plasmids and chromosomes are associated to individual genomes, we know which of these elements co-occur. We retrieved 1943 complete phages genomes from NCBI RefSeq in November 2016. The lifestyle of these phages was predicted using PHACTS v.0.3^23^. Predictions were considered as confident if the average probability score of the predicted lifestyle was at least two standard deviations (SD) away from the average probability score of the other lifestyle, as recommended by the authors (who claim a precision rate of 99% with this parameter). Using these criteria, we classified 54% of the phages into 420 virulent and 629 temperate phages.

### Prophages detection

We detected 9927 putative prophages in the bacterial genomes using PhageFinder v.4.6 as in^24^ (http://phage-finder.sourceforge.net/). The elements smaller than 10 kb and with >25% of the predicted genes belonging to ISs were removed. Such elements identified by Phage Finder may be prophage remnants or erroneous assignments.

### CRISPRs and Cas clusters detection

Cas clusters were detected with MacSyFinder v.1.0.2^25^, within CRISPR-CasFinder v.0.9, which classed them in subtypes following ^8^. They correspond to the set of minimal genes that is thought to provide a functional system for a given subtype. While this work was being finished, a new version of CRISPR-CasFinder (v.1.0) was released ^26^. We compared the results provided by the two versions (Supplementary Figure 1) and found them to be almost identical. Version 1.0 only detected 38 additional clusters (1% of the dataset). The program is available at https://crisprcas.i2bc.paris-saclay.fr/CrisprCasFinder/Index. All results are reported in Supplementary Table 1.

**Figure 1:**
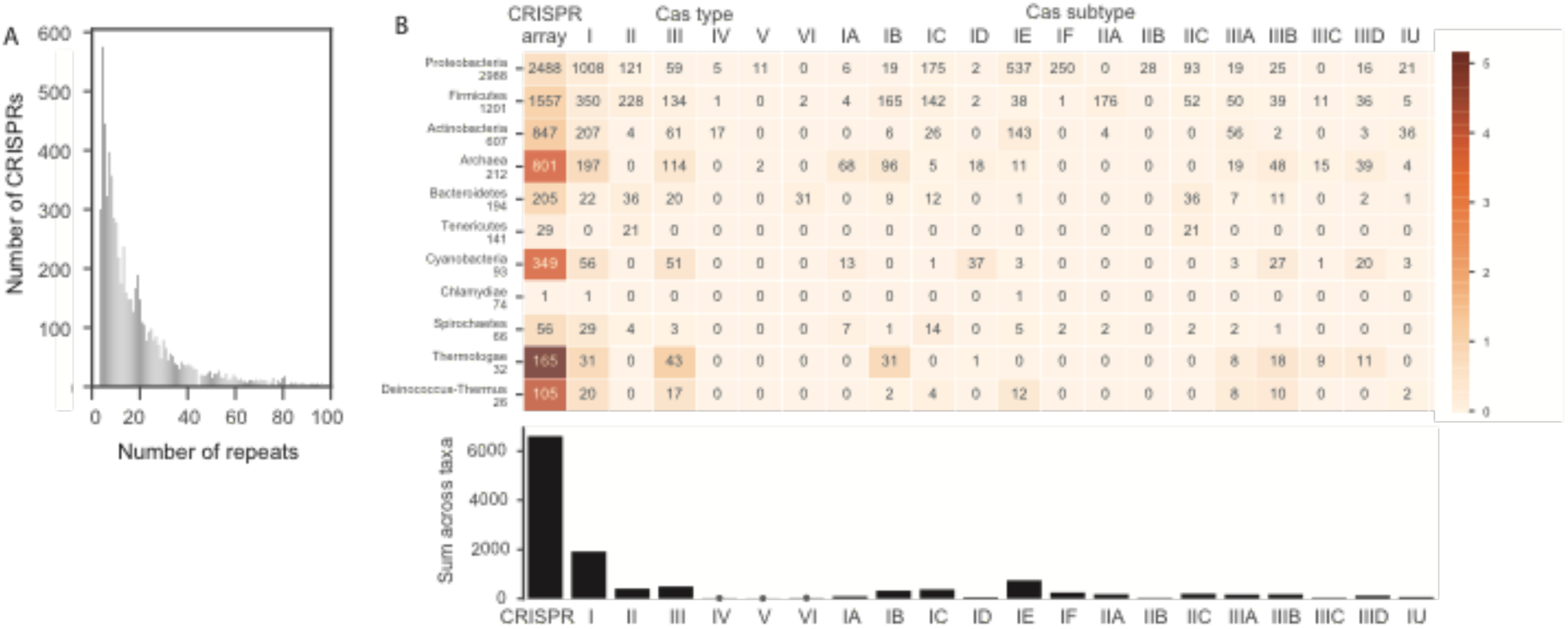
Distribution of CRISPR arrays and clusters of *cas* genes in the genomes of Prokaryotes. **A.** Histogram of the number of repeats in CRISPRs (histogram truncated at 100, because higher values are very rare, maximum is 589). **B.** Distribution per clade (on the top panel only clades with more than 25 genomes are indicated). The cells indicate the number of systems detected in the clade, and the colour of the cell is proportional to the average frequency per genome (the darker, the more frequent, see scale). The bottom panel shows the total number of elements detected in the dataset.

We detected CRISPRs using the CRISPR Recognition Tool v.1.2 (CRT)^27^ with the default parameters except for the maximum length of a CRISPR’s repeated region (–maxRL) which was set to 50. To limit the number of false positives, we calculated the coefficient of variation of the length of the spacers within each CRISPR. This coefficient is expected to be low, as spacers are integrated through mechanisms that ensure a specific size^28^. To define a threshold above which detected arrays would be considered as false (*i.e.*, detected repeats would not be CRISPR), we computed the coefficient of variation of length of spacers of CRISPRs close to *cas* clusters (here 10 kb). We restricted this analysis to this dataset because CRISPR neighbouring *cas* clusters are unlikely to be false (Supplementary Figure 2). The coefficient of variation observed for the length of these spacers was rarely larger than 0.19 and almost never larger than 0.28. We therefore chose to remove from the dataset the CRISPR arrays with a coefficient of variation superior to 0.28. This step removed 318 arrays representing only 4.4% of the dataset. All results are reported in Supplementary Table 2.

**Figure 2:**
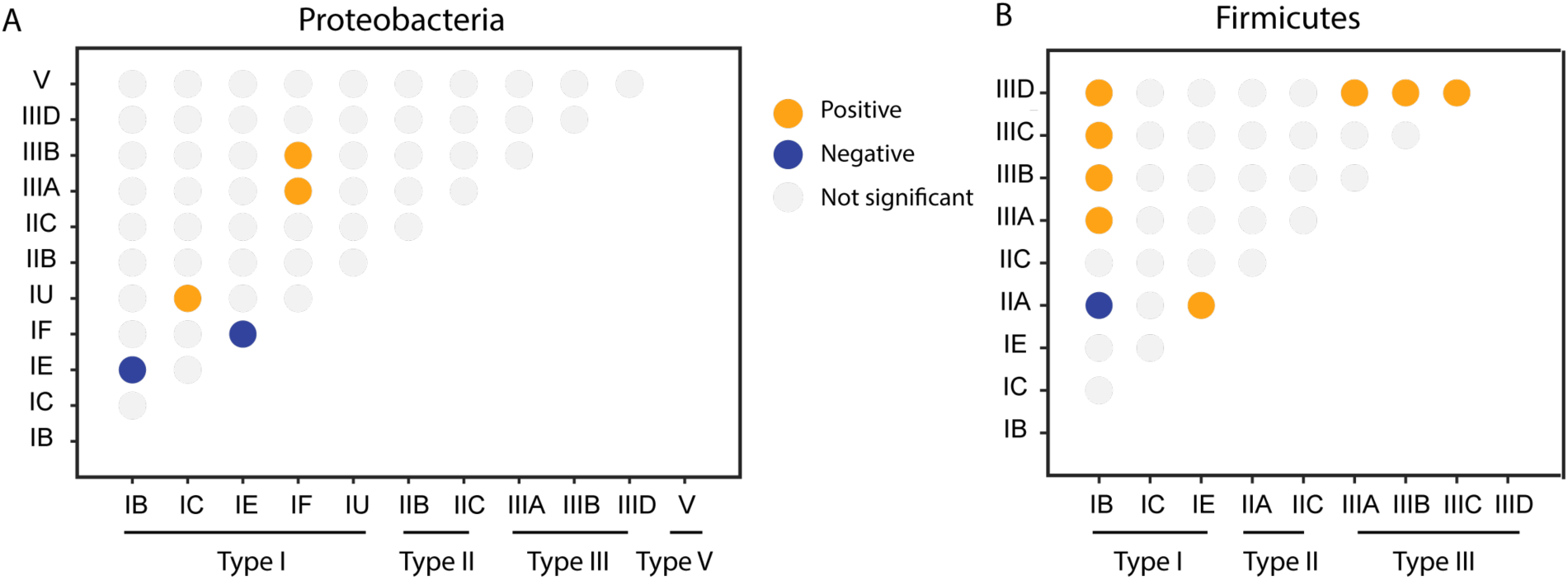
Significant associations between Cas subtypes in the same genome in Proteobacteria (A) and Firmicutes (B). Each circle corresponds to the association between two subtypes. Associations are represented in grey (not significant), blue (negative), and orange (positive). Only sub-types with systems present in more than 1% of the genomes in the clade are represented (the others never have significant statistics).

### Linking CRISPRs and Cas clusters

In order to be able to associate CRISPRs with cognate *cas* clusters, we calculated the minimal circular distance between an array and a cluster. When CRISPR and *cas* clusters were at distances lower than 20 kb, they were put together in one single CRISPR-Cas locus. The clustering was done by transitivity, if other elements, CRISPRs or *cas* clusters, were less than 20 kb away from the locus, they were also assigned to the locus. Hence, a CRISPR-Cas locus is a region in the chromosome containing at least one *cas* cluster and one CRISPR, where the elements are never more than 20 kb from the closest element. A specific case occurs when one or several CRISPR are close to one single cas cluster in a CRISPR-Cas locus. In this case, we assigned a subtype to the CRISPR array (the one of the single *cas* cluster). Subtypes could not be assigned with this method to CRISPRs outside CRISPR-Cas loci and to CRISPRs in loci containing more than one subtype of *cas* clusters.

To assign a type to the CRISPRs outside CRISPR-Cas loci, we searched to similarities between their repeats and those of CRISPRs that could be types with the method described above. We used the information on the CRISPR subtypes, taken from the CRISPR-Cas loci with a single Cas subtype, to build a databank of 3324 unique repeats (direct and reverse complement sequences) that we could confidently assign to specific Cas subtypes. This was then used to type the other repeats (those in the remaining CRISPRs). For this, we quantified the sequence similarity between every pair of CRISPR repeats using a global alignment with no gap end penalty and equal gap creation and extension penalties (−3) using the module pairwise2 from Biopython (function align.globalxs). As a result of this procedure, we obtained a table where each pair of repeats is associated with a continuous variable indicating the sequence identity between the two repeats and with a binary variable indicating if the two repeats are from the same Cas subtype. We used this data to make a logistic regression between the identity score of the best hit and the categorical variable assigning the subtype prediction. We used a ROC curve to choose a threshold with a high True Positive Rate (80%). At this threshold, if the best hit among the repeats of an unknown type CRISPR matches a repeat of a CRISPR of a given subtype with more than 72% identity, the first CRISPR is classed as of the same subtype as the second.

Having defined a minimal sequence identity to class a repeat into a Cas subtype, we used it to assign subtypes to the CRISPRs. For each array, we quantified the sequence identity of its repeats with all repeats of the typed CRISPR repeats. We used a global alignment with no gap end penalty and equal gap creation and extension penalties (−3) using the module pairwise2 from Biopython (function align.globalxs). We took the best hit among those scores. If the identity score was higher than 72%, we classed the array of repeats to the subtype of the best hit.

### Phylogenetic analyses

We identified the families of orthologous proteins present in more than 90% of the genomes (when larger than 1 Mb) of two phyla: Firmicutes (1189 genomes), and Proteobacteria (2897 genomes). The genomes were obtained from GenBank’s RefSeq dataset as indicated above. The orthologs were identified as reciprocal best hits using end-gap free global alignment, between the proteome of a pivot and each of the other strain’s proteomes (as in ^29^). *Escherichia coli* K12 MG1655 and *Bacillus subtilis* str.168 were used as a pivot for each clade. Hits with less than 37% similarity in amino acid sequence and more than 20% difference in protein length were discarded. The persistent genome of each clade was defined as the intersection of pairwise lists of orthologs that were present in at least 90% of the genomes representing 411 protein families for Firmicutes and 341 for Proteobacteria.

We inferred phylogenetic trees for each phyla from the concatenate of the multiple alignments of the persistent genes obtained with MAFFT v.7.205 (with default options)^30^. Alignments were purged of poorly informative sites using BMGE v1.12 (with default options)^31^. Missing genes were replaced by stretches of “-” in each multiple alignment. Adding a small number of “-” has little impact on phylogeny reconstruction^32^. The trees of the phyla were computed with FastTree version 2.1 using the LG model^33^, which had lower AIC than the alternative WAG model in both cases. We made 100 bootstrap trees using phylip’s SEQBOOT^34^ to generate resampled alignments which were given as input to FastTree (options –n –intree1).

We tested the association between types of Cas systems in two steps. First, to lower the amount of computational load, we assumed that bacteria were phylogenetically independent. We build a contingency table for each pair of types of Cas. We selected the pairs for which the hypothesis of independence was rejected (p<0.05, Fisher exact test). These pairs were then re-analysed to account for phylogenetic dependency using BayesTraits v.3^35^, and the abovementioned phylogenetic tree. We ran the likelihood estimation of two models: independent or dependent evolution of two traits. We performed a Likelihood-ratio test to compare the two models for each of the 100 trees. We validated an association if the median of the p-values of the 100 Likelihood-ratio test was inferior to 0.01. We did not use the Bayesian version of BayesTraits because it could not handle the very large size of our dataset.

## Results

### Epistatic interactions between *cas* clusters

We identified 6949 CRISPRs and 3251 clusters of *cas* genes in fully sequenced bacterial (5563) and archaeal genomes (212) (Supplementary table 1). The size of CRISPRs varies widely, from a minimum of three repeats (minimal detection threshold) to a maximum of 589 in the Proteobacteria *Haliangium ochraceum* DSM 14365 (Supplementary Table 4). Most arrays are small, with 19% of them containing between three and five repeats (Figure 1.A). CRISPRs were found in 47%, and Cas clusters in 42% of the bacterial genomes. The distribution of Cas types is very heterogeneous. The Type I Cas systems are by far the most frequent (present in 30% of all genomes) followed by type II (7%) and type III (8%) (Figure 1.B). All of them are present in different phyla. The types IV, V and VI are extremely rare in the current genome database – they were found in less than 30 genomes – and are exclusive of a few clades (Proteobacteria, Actinobacteria, Bacteroidetes). Some subtypes are present in many clades – I-B, I-C, II-C, III-A, III-B, III-D – while others are clade specific, e.g., subtype I-D is mostly found in Cyanobacteria, II-A in Firmicutes and II-B in Proteobacteria (Figure 1.B). The relative abundances of CRISPR and Cas subtypes are close to those reported in previous studies^8,25^. Overall, the number of systems per genome does not show systematic variations with genome size, except that small (<1 Mb) genomes rarely encode these systems (Supplementary Figure 3).

**Figure 3:**
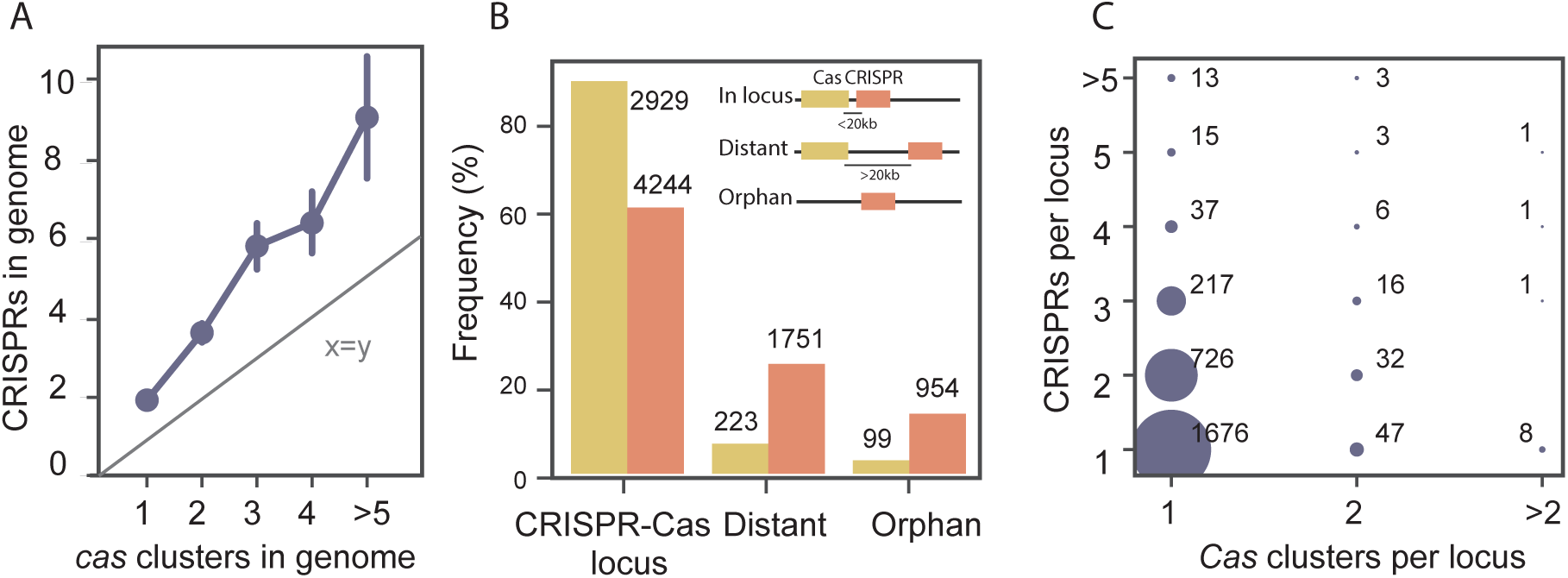
Organization of CRISPR-Cas loci. **A.** Number of CRISPR arrays in function of the number of *cas* clusters in bacterial genomes (mean, CI 95%). The straight line indicates the identity (number of CRISPRs equal to the number of Cas clusters) **B.** Frequency of CRISPRs and *cas* clusters in terms of their genetic context. Loci were classed as complete CRISPR-Cas loci when they included at least one CRISPR and one *cas* cluster, “distant” when the element (CRISPR or *cas* cluster) is more than 20 kb from the closest cognate element, and “orphan” when the cognate element is absent from the genome. **C.** Quantification of the different organizations of CRISPR-Cas loci.

Most genomes lack *cas* clusters, but some others have multiple clusters. To understand if epistatic interactions between different systems could explain the co-occurrence of these multiple *cas* clusters, we analysed the co-occurrence of all pairs of Cas types. We used Bayestraits to integrate the information of the phylogenetic structure in the evaluation of these associations ^35^. Since phylogenetic inference of all the prokaryotes is very inaccurate, we restricted our analysis to Firmicutes and Proteobacteria, the two clades with more genomes in our dataset (75% of the total). We inferred 100 phylogenetic trees for these clades (to account for uncertainty in phylogenetic inference), and used them to test the associations between every pair of systems (Figure 2.A and 2.B). We found some cases of unexpectedly low co-occurrence of Cas subtypes. In Proteobacteria, subtype I-E is negatively associated to both I-B (never observed) and I-F (observed 17 times, expected 45). In Firmicutes, only subtypes II-A and I-B co-occur less than expected. Negative co-occurrences between Cas subtypes could be explained by counter-selection following negative epistatic interactions between systems, or by functional redundancy leading to the loss by drift of one of them. Types I-E and I-F are very similar and occur in several genomes, suggesting that their joint presence in a genome is not deleterious in certain genetic backgrounds. In contrast, type I-B and I-E are never observed together, even if the expected number of co-occurrences is low (4). Subtypes II-A and I-B are from the two different large Cas classes and are less likely to be redundant. The lower than expected co-occurrence of these systems suggests incompatibility between their machineries.

Higher than expected co-occurrences of Cas subtypes were abundant. This may indicate selection for certain combinations of adaptive immunity mechanisms. In Proteobacteria, we observed positive associations between subtype I-U with I-C (but only 4 occurrences are observed). The deviations from the expected values are higher for the co-occurrence between subtype I-F and III-A/B systems (12 observed, 4 expected). This is in agreement with experimental work showing that in *Marinomonas mediterranea* (a Proteobacteria), there is synergy between the action of type I-F and type III-B systems^21^. In Firmicutes, the network of significantly high co-occurrences is more structured and shows stronger deviations from the expected random distribution than in Proteobacteria. Notably systems I-B co-occur more than expected with all type III systems (72 observed, 13 expected), subtype I-E co-occurs more than expected with subtype II-A (15 observed, 5 expected), and subtype III-D is more often present in genomes encoding any of the other type III systems than expected (15 observed, 3 expected). Overall, there is a clear excess of positive co-occurrences of type I and type III systems, relative to other combinations.

### Many CRISPR-Cas loci are complex

We observed that many genomes with a *cas* cluster had more than one cluster (27%), and most genomes with a CRISPR had more than one array (60%). This challenges the canonical view of the organization of a CRISPR-Cas locus as the association of one CRISPR with one *cas* cluster. Accordingly, genomes encode more CRISPRs than *cas* clusters and, in genomes encoding several of the latter, the number of CRISPR arrays grows faster than the number of *cas* clusters (Figure 3.A). This suggests that increasing the number of CRISPRs per *cas* cluster is beneficial in bacteria were CRISPR-Cas immunity plays an important role. This fits theoretical works suggesting that the presence of multiple CRISPR loci in a genome can be adaptive if they have different spacer turnover rates^36^.

To shed some light on the multiplicity of CRISPRs and *cas* clusters, we must first solve the problem of how to associate them in CRISPR-Cas loci. This problem is trivial when there is one single *cas* cluster and a contiguous single CRISPR, but not when there are multiple *cas* clusters or CRISPRs. The distributions of the distances between *cas* clusters and the closest CRISPR and between a CRISPR and the closest *cas* cluster (they are not identical because there are more CRISPR than *cas* clusters), reveal three groups: very close elements (<700 bp), intermediate (between 700 bp and 20 kb) and distant (>20 kb apart) (Supplementary Figure 4.A and B). The careful analysis of the “intermediate” group showed that the sequences intervening between the CRISPR and the *cas* cluster were often either other CRISPRs or genes that might be associated with the *cas* clusters. The latter were not annotated by our pipeline because we focus on the most conserved genes^37^. The probability of finding pairs of elements less than 20 kb by chance is much lower than that observed in genomes. Based on these arguments, we defined a *CRISPR-Cas locus* as a region in the genome containing at least one *cas* cluster and one CRISPR, and eventually other such elements when two consecutive elements are spaced by less than 20 kb (clustered by transitivity, Supplementary Figure 4.C). Hence, multiple *cas* clusters and CRISPRs can be part of the same locus. The elements not included in CRISPR-Cas loci were classed in two categories. *Distant* elements are CRISPRs or *cas* clusters more than 20 kb away from the closest cognate element. *Orphan* elements are those present in genomes lacking the cognate element (*i.e.*, CRISPRs in genomes without *cas* clusters and vice-versa). Using this classification, the vast majority of *cas* clusters (90%), and a small majority of CRISPRs (61%) are part of CRISPR-Cas loci. Around 25% of the CRISPRs are distant and 14% are orphans (Figure 3.B). Hence, there is an asymmetry in the genetic organization of the components of these systems: *cas* clusters are much more often co-localized with CRISPRs than the latter are co-localized with *cas* clusters.

**Figure 4:**
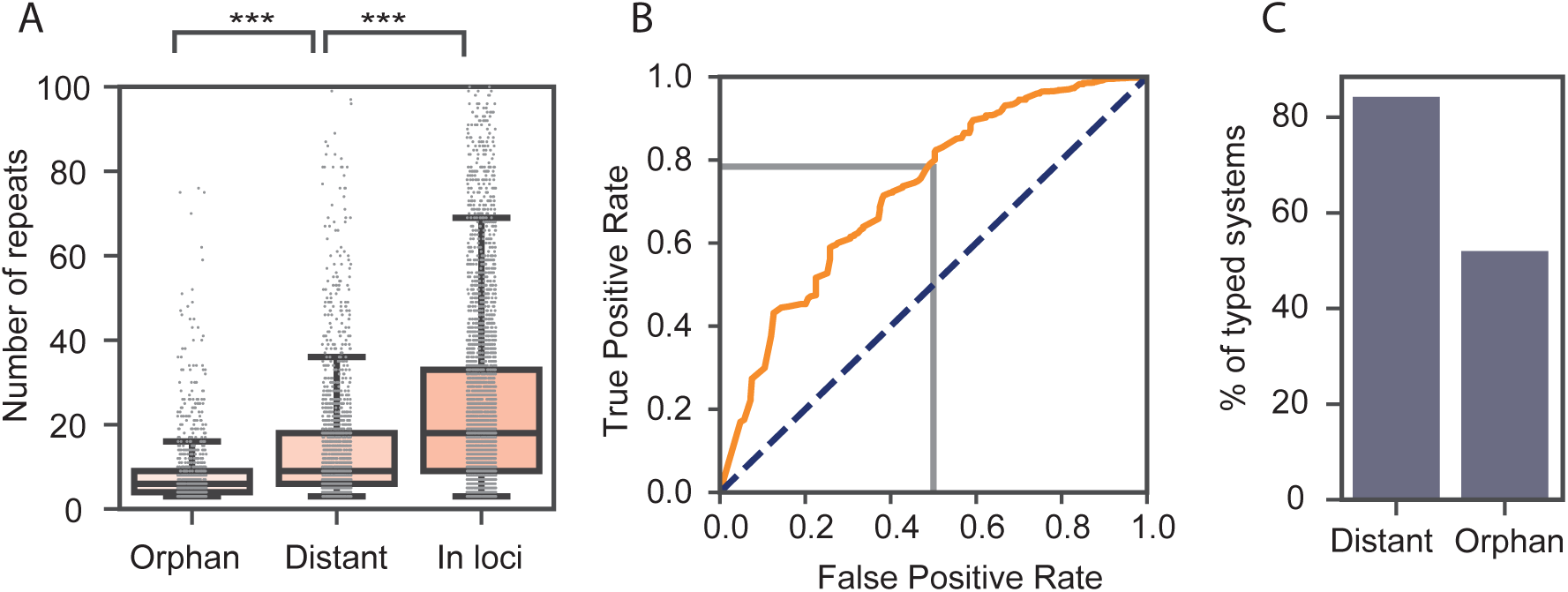
Characterization of CRISPRs according to their association with *cas* clusters. **A.** Number of repeats in CRISPRs in function of their distance to *cas* clusters (Tukey HSD, all pairs, P<0.001). **B.** ROC curve (orange) of the results of the study using logistic regression to predict the subtype of Cas systems for the best hit of the set of repeats of a CRISPR. In grey, the threshold chosen to assign subtype to unknown arrays (72% identity). **C.** Percentage of orphan and distant arrays with sub-type assignment.

We classified CRISPR-Cas loci in function of the number of CRISPRs and *cas* clusters (Figure 3.C). The canonical CRISPR-Cas system – a locus with one CRISPR and one *cas* cluster - represents slightly more than half of all loci (60%). More than a fourth of all loci have one *cas* cluster and two CRISPRs. Many other combinations are observed, even if they are rarer. Among these, we observed that 4% of the loci encode more than one *cas* cluster. This shows that the organization of the loci can be much more complex than the prototypical one *cas* to one CRISPR textbook example.

### Most distant CRISPR systems could function *in trans*

The above classification allows to investigate more closely the association between CRISPRs and *cas* clusters in CRISPR-Cas loci. Within CRISPR-Cas loci with a single *cas* cluster, the size of the CRISPRs depends on the subtype of the *cas* cluster (Supplementary Table 4, ANOVA, P<0.001). Type IV, V, VI and subtype II-A tend to have short CRISPRs (<20 repeats on average). On the other hand, subtype I-A, I-B, I-D have the longest CRISPRs with more than 40 repeats on average. CRISPRs outside CRISPR-Cas loci are different. Orphan CRISPR arrays are smaller (9 repeats on average) than distant arrays (16), which are smaller than arrays within CRISPR-Cas loci (28, Figure 4.A). Consequently, the presence, proximity and subtype of *cas* clusters impact the number of repeats in CRISPRs.

We put forward the hypothesis that CRISPRs distant from CRISPR-Cas loci could be mobilized by the latter for immune defence. If true, their repeats should match those of the *cas* clusters *in trans*. To test this hypothesis, we typed the CRISPRs outside CRISPR-Cas loci and checked if they matched the subtype of CRISPR-Cas systems in the genome. We typed these CRISPRs using the information on the best hit of the repeats of the CRISPR to a databank of 3602 unique repeats that could be unambiguously assigned to a Cas subtype (because they were taken from CRISPRs of CRISPR-Cas loci with one single *cas* cluster, Supplementary Table 2). We used a logistic regression to set the identity score threshold above which a best hit could be reliably associated with a Cas subtype. We chose a threshold that corresponds to an identity score of 72% (Figure 4.B, Supplementary Figure 5.A). The analysis of the original dataset shows 2773 correct and 736 incorrect assignments (accuracy of 79%). This method allowed the assignment of subtypes to 50% of the orphan arrays and 85% of the distant arrays (Figure 4.C). The different levels of success in typing the two classes of CRISPR may be explained by the presence of spurious CRISPR arrays in the orphan dataset^38^. Accordingly, untyped orphan CRISPRs are shorter on average (7) than the typed ones (11 repeats, Supplementary Figure 5.B, Mann Whitney, P<0.001). This suggests that many of the untyped CRISPRs might be either false positives or elements ongoing genetic degradation (which is presumably facilitated by the lack of a cognate *cas* cluster in the genome). The analysis of distant CRISPRs revealed that 60% of them had repeats of the same subtype as the *cas* cluster present in the genome *in trans*. The relatively high number (40%) of non-matching repeats changed only slightly (43%) when the analysis was restricted to arrays with a number of repeats higher than 5 (Supplementary Table S3). Hence, the majority of CRISPRs distant from *cas* clusters have repeats matching these clusters. This suggests that such CRISPRs could be used by the latter for immune defence.

**Figure 5:**
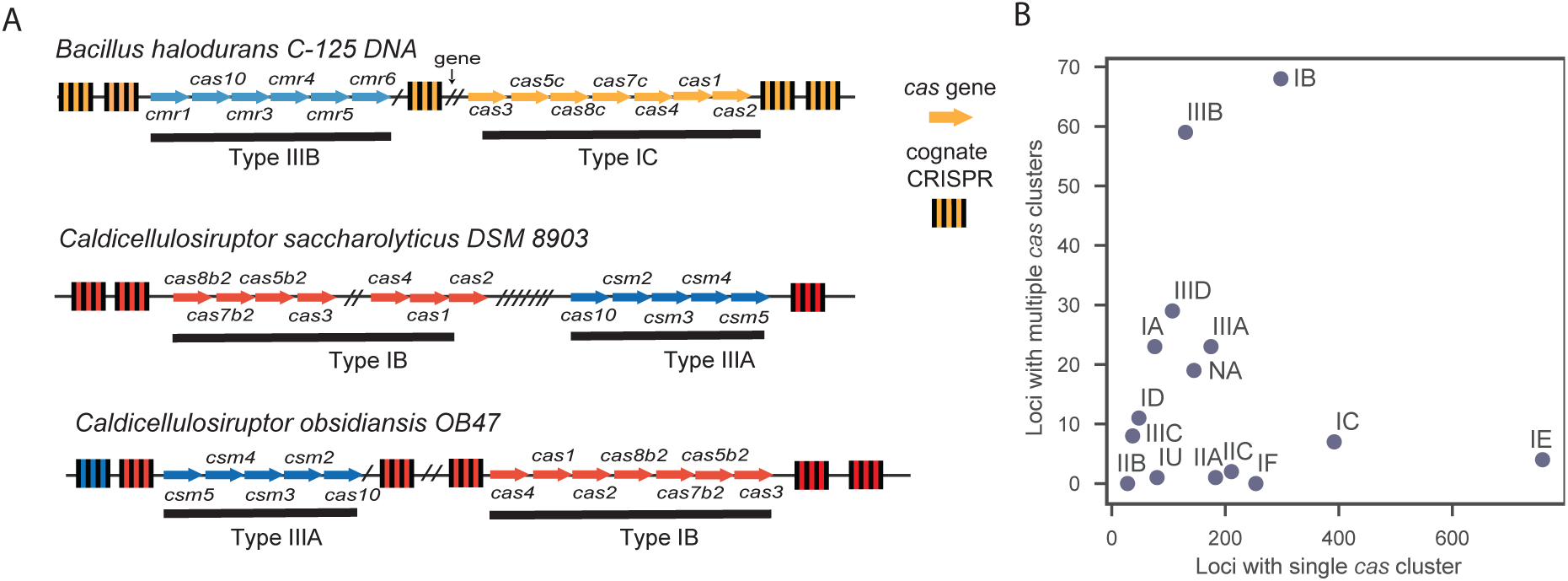
Association between Cas clusters. **A.** Examples of complex CRISPR-Cas loci found in Bacteria. Arrows represent *cas* genes and cas clusters are coloured by subtypes. Genes that were not identified as *cas* genes were omitted and replaced by a slash (/). CRISPRs colours match the Cas subtype to which their repeats were assigned. Grey indicates that no subtype could be assigned. **B.** Number of loci with a given Cas subtype found in simple or complex loci.

### Complex loci have unique adaptation and multiple interference mechanisms

The analysis of co-occurrence of *cas* clusters and the ability to type CRISPRs using the sequence of their repeats paves the way to study in detail the complex CRISPR-Cas loci, especially those with more than one Cas subtype. Analysis of these loci shows that different *cas* clusters are often clearly separated by other elements, such as CRISPRs (Figure 5.A). Other genes that were not annotated by our procedure are also found between *cas* clusters or between pairs of CRISPRs. Some of them have previously been proposed to be associated with CRISPR-Cas systems^37^. Some Cas subtypes are more likely to co-occur in a locus than others (Figure 5.B). For example, type II systems rarely co-occur with other systems in the same locus. This is also the case of most type I systems, with notable exception of type I-B and at a lesser extent I-A. Type III elements are much more likely to be in complex CRISPR-Cas loci. In particular, subtypes I-B and III-B were very often found together (27% of all complex loci). This fits the previous observation of positive genome-wide associations between subtype I-B and type III systems in Firmicutes (Figure 2.B). Interestingly, although few *cas* clusters in genomes could not be typed (less than 10%), they are often found in complex CRISPR-Cas loci. This could be explained by functional interaction between *cas* clusters leading to the loss of certain *cas* genes that render the specific system hard to type.

Complex CRISPR-Cas loci often have several CRISPRs and *cas* clusters. We wondered if there were preferential associations between the two. The CRISPRs in complex loci with multiple *cas* clusters have identical repeats in 59% of the cases, and repeats are more than 80% identical in 48% of the remaining cases. Hence, there is less heterogeneity among CRISPRs than expected given that these loci have different Cas subtypes. This is in line with our observation that these loci have one single pair of *cas1*-*cas2* genes, the key genes involved in adaptation, in the majority of cases (86% have only one *cas1*). To further test the hypothesis that loci with multiple *cas* clusters often share one single adaptation module, we searched to identify the Cas subtype associated with the CRISPRs. In 101 out 118 loci, all the CRISPRs that could be typed in a locus were from the same subtype. In particular, clusters with Cas of types I-B and types III had repeats classed as I-B 84% of the times (31 out of 37 cases) showing that these loci tend to have the adaptation module of type I-B systems. These results suggest that complex loci use multiple mechanisms for interference and a single, often type I-like, mechanism for adaptation.

### CRISPRs in mobile elements match chromosomal Cas systems

The ICP1 phage of Vibrio cholerae carries a CRISPR-Cas system to subvert its host immune defences^19^. Following on this study, we wished to assess the frequency with which CRISPR-Cas systems occur in phages and if there are significant associations between these systems and those of the host. We quantified the occurrence of CRISPR-Cas systems in the 1943 phages of RefSeq. Using PHACTS, we could characterize the lifestyle of more than half of the phages: 40% are virulent and 60% are temperate. We found only one *cas* cluster and five CRISPRs in these genomes (Figure 6). We then analysed the sequences of 9926 prophages to search if temperate phages that successfully infected a host were more likely to encode CRISPR-Cas systems than the other phages. We only detected two complete CRISPR-Cas systems, six *cas* clusters and 33 CRISPRs. These values are very similar to those identified in the phage dataset, once the different number of elements is accounted for. We conclude that, within the limits given by the size and diversity of current genome databases, CRISPR-Cas systems are extremely rare in phages. Their frequency in prophages is not significantly different from that of the average phage.

**Figure 6:**
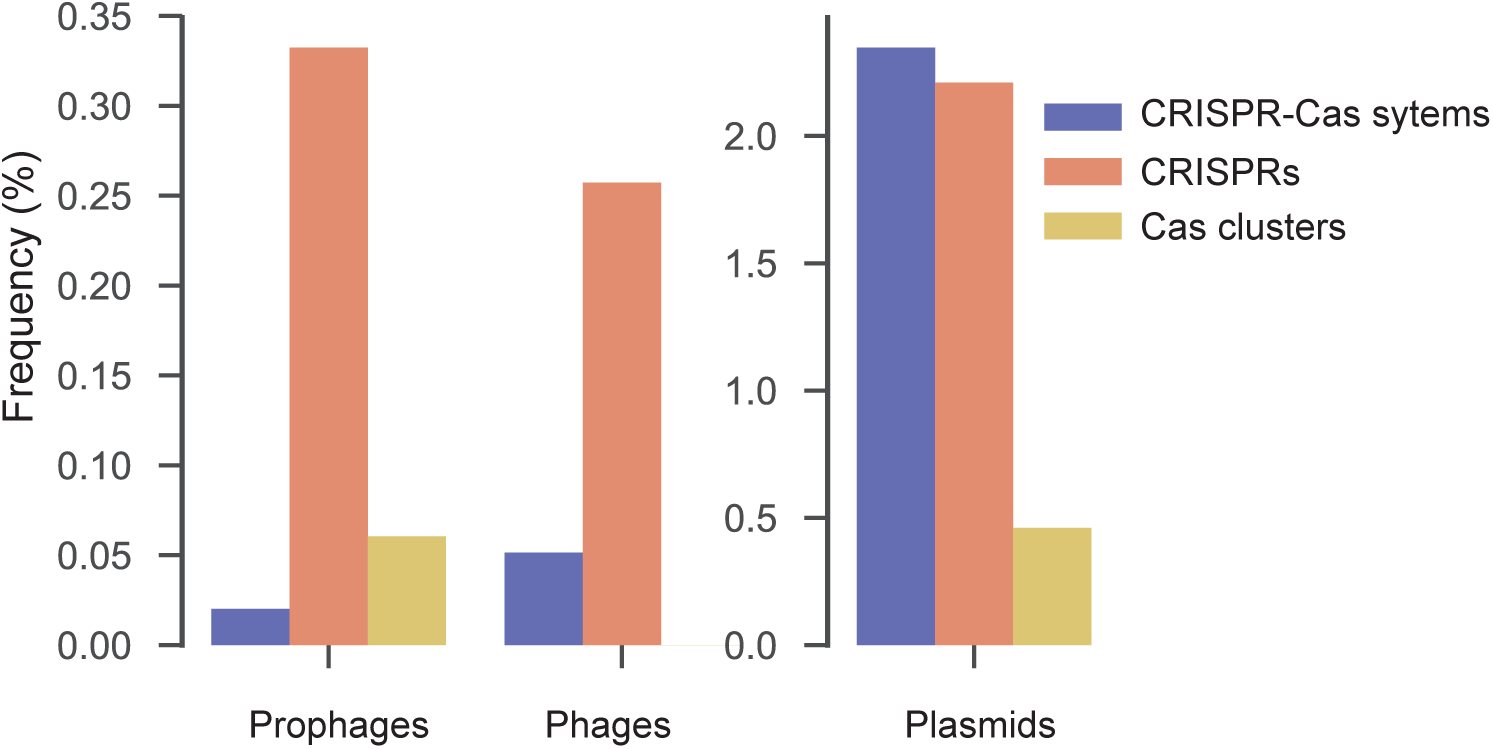
Frequency of CRISPR-Cas systems in prophages, phages and plasmids.

We then turned our attention to plasmids. We searched for CRISPR-Cas systems among 4335 plasmids that were sequenced in the whole-genome projects analysed in this study. This means that we know the chromosome of the bacterial host of every plasmid in the database. We found 112 complete systems and 101 CRISPRs in plasmids devoid of *cas* clusters (Figure 6). Plasmid CRISPRs have on average 15 repeats, significantly less than chromosomal arrays (22 repeats) (Supplementary Figure 6.A Mann Whitney, P<0.001). The relative abundance of subtypes is also significantly different on plasmids and chromosomes (Supplementary Figure 6.C). In particular, no plasmids encode type II-A CRISPR-Cas systems while type IV systems are encoded almost exclusively on plasmids. Plasmids with CRISPR arrays and encoding *cas* clusters were larger (1.5 times when only encoding a CRISPR and 2.5 times when encoding a Cas) than the other plasmids (Supplementary Figure 3.C). These results show that plasmids are much more likely to encode CRISPR and especially CRISPR-Cas loci than phages, even if this concerns less than 5% of all plasmids. The differences observed between the distributions of Cas subtypes in plasmids and chromosomes suggests that plasmid CRISPR-Cas systems are not just a random sample of chromosome systems. Instead, they may reflect selection for systems influencing the interactions between the plasmid and its host.

The large number of plasmids carrying CRISPRs but lacking *cas* clusters suggests the existence of interactions between plasmid-associated CRISPRs and the chromosomal-encoded Cas proteins. Almost half (48%) of the genomes with CRISPRs in plasmids (but no *cas* clusters) have chromosomal *cas* clusters (Supplementary Figure 7.A). We typed these CRISPRs and then tested if they matched the subtype of *cas* clusters found on the chromosome. We assigned a subtype to 27 of these plasmid CRISPR arrays, and in 16 cases these matched the type of the chromosomal *cas* clusters (Supplementary Figure 7.B). We tested the significance of this result by simulating the expected number of matches between plasmid and chromosomal subtypes. In 1000 simulations, the average number of matches was 6.4 and the highest number was 13 (Supplementary Figure 7.C). While the number of observations is low, these results suggests that when plasmids with CRISPRs, but no *cas* clusters, are in genomes with *cas* clusters, the array is more likely to be classed in that precise Cas subtype than expected by chance.

## Discussion

We detected and analysed thousands of CRISPRs and *cas* clusters in fully sequenced bacterial and archaeal genomes, the complete content of the NCBI RefSeq database at the moment of the start of our study. Hence, there was no bias in our analysis of the data, apart from the bias of the database itself, which is known to over-represent cultivable Bacteria over other Prokaryotes. It is possible that extremely rare systems in our dataset – types IV, V, and VI- are more frequent in poorly sampled clades. However, it should be noted that these three systems tended to be overrepresented in phyla that are well sampled in our database (Proteobacteria, Actinobacteria, and Bacteroidetes). Further work and much broader samples will be needed to understand if these systems are rare and why this is so. In this study, their low frequency resulted in no significant association with other systems. Our analysis also revealed that 19% of CRISPRs have less than five repeats. Half of these CRISPRs with three to five spacers were in CRISPR-Cas loci, but part of the other small CRISPRs might be false positives. To control for their impact on our results - when necessary – we made replicates of the analyses using only CRISPRs with five or more repeats. These analyses resulted in qualitatively similar conclusions (Table S3). We used CasFinder to identify and type *cas* clusters, which was previously shown to provide accurate classifications^26^. As our detection takes into account the architectures and signature proteins of *cas* clusters, it provides a robust subtype assignment compared to a previous study where subtypes were only inferred from Cas1^13^. To associate the CRISPRs outside CRISPR-Cas loci to Cas types we used the sequence similarity to a database of known repeats. This part of our method might gain from the diversification of the genome database, since rare CRISPR-Cas systems are under-represented. A larger reference database might also allow to define subtype-specific sequence identity thresholds. Such developments could pave the way to understand if our inability to type half of the orphan CRISPRs (because they lack high sequence similarity to known repeats) is due to the lack of a sufficiently comprehensive repeat database or to the rapid evolution by drift of orphan arrays in the absence of *cas* clusters.

The goal of this study was to understand the association between Cas and CRISPRs, but this led to methodological developments in CRISPR typing that could be useful in metagenomic studies, where most of the detected elements are orphan because most contigs are very small. We show in supplementary material a proof of concept for the classification of CRISPRs from metagenomics data (Supplementary Text1, Supplementary figure 5.C).

Consistent with previous analyses^8,13^, we observe that most bacteria lack CRISPR-Cas systems. This is puzzling, because most species have mobile genetic elements against which these systems might presumably provide protection. For example, half of the genomes in the database are lysogens^24^, and many have plasmids (this work). If CRISPR-Cas are universally efficient immune systems, why is it that most bacteria lack them? And why is that many of the remaining bacteria have small CRISPRs? This cannot be caused by phylogenetic inertia, *i.e.* the fact that certain lineages have not developed such systems, since CRISPR-Cas systems are frequently transferred across phyla^14,15,39,40^. Instead, it has been proposed that the deleterious effects of CRISPR-Cas systems on the host, either by spacers targeting the chromosome^41^, or by interfering with DNA repair functions^42^ could explain the relative paucity of CRISPR-Cas systems across Bacteria. These processes may also explain why so many CRISPRs contain so few spacers: they could result from decaying inactive CRISPR-Cas systems. The size of the CRISPRs may also be affected by the balance of the rates of acquisition and loss of the spacers. Experimental observations on primed adaptation (acquisition of spacers from a mobile elements already targeted by a spacer in the CRISPR)^43^ and some mathematical models predict selective sweeps of lineages with CRISPR-Cas systems effective in providing immunity against phages present in the community^44^. CRISPRs could thus acquire several spacers within a short time-frame, rapidly increasing in size, whereas the loss of old spacers is more gradual^45,46^. As a result, short CRISPRs could result from the gradual shrinkage of CRISPR arrays that have not undergone recent acquisition – selection events. The small CRISPRs found in this and previous studies suggest that CRISPRs target a relatively small number of mobile genetic elements in most individual genomes.

Orphan or distant CRISPRs represent 40% of all the CRISPRs. Long gaps in the activity of CRISPR-Cas systems might also explain the abundance of CRISPRs without a neighboring *cas* cluster. While the system is not being used, and therefore is not adaptive, *cas* clusters may be lost in a neutral manner. Similar neutral processes may take place when CRISPR-Cas systems are acquired by a host that is incapable of expressing the *cas* genes. Such processes of genetic erosion should be accelerated when the systems turn out to be deleterious in the novel genetic background. Since it takes *cas* genes to provide a function to CRISPR spacers, the former are likely to be lost first, resulting in many CRISPRs without cognate *cas* clusters. Finally, mobile elements drive most of the horizontal gene transfer across species and may also increase the frequency of isolated CRISPRs. We have shown that mobile elements have more (in phages), or almost as many (in plasmids) CRISPRs than CRISPR-Cas loci. In plasmids, we could show that these CRISPRs match the chromosomal *cas* clusters more often than expected, suggesting that they evolve to interact with the host systems. Integration of these mobile elements in the genome leads to a further excess of CRISPRs relative to *cas* clusters. Finally, chromosomal distant CRISPRs could originate from CRISPR-Cas loci by genome rearrangements, even if these are rarely fixed^47^. In this case, the CRISPRs distant from the *cas* clusters would be compatible with them, as often observed in our data. The smaller size of these CRISPRs could result from reduced efficiency at incorporating spacers when arrays are distant from *cas* clusters or less efficient selection for the spacers if expression of these arrays is weaker than those located next to *cas* clusters.

Little was known about the frequency of CRISPR-Cas in phages and plasmids, in spite of the previous studies describing their existence and relevance^16–19^. A recent preprint suggests that CRISPR-Cas may be particularly abundant in the phages of very large genomes^48^. Yet, we show that CRISPR-Cas are almost never carried by the phages available in the genome databases. This does not invalidate the previous results showing that CRISPR-Cas carried by phages may provide the latter with a mechanism to escape host innate immunity^19^. Yet, if the current phage database is representative of the natural diversity of these elements, such mechanisms are rarely used by phages and may be specific of a few families. Interestingly, the much higher frequency of plasmids carrying CRISPR-Cas and especially CRISPRs compatible with the chromosomal *cas* clusters opens the possibility for plasmids to manipulate the host immunity by using the host Cas proteins and their own CRISPRs. The differences in terms of the distributions of Cas subtypes in plasmids and chromosomes reinforces this hypothesis, because it suggests that plasmid systems are not just random samples of chromosomal systems. This is particularly striking in the case of the type IV systems, previously reported in plasmids^8^, that we show are almost exclusively encoded on these mobile elements. As these systems do not encode Cas1 and Cas2, the main proteins for adaptation^8^, it is not known how they acquire new spacers. It is tempting to speculate that type IV CRISPR are able to use the adaptation machinery of the host’s CRISPR-Cas systems, in a way resembling the CRISPRs in plasmids matching chromosomal Cas systems, and that of type III systems in complex loci that share the adaptation machineries of type I systems. If true, and given the vast over-representation of type IV systems in plasmids, it suggests that these systems may have evolved as specialists in subverting chromosomal systems. This is consistent with the observation that these systems frequently lack not only the adaptation machinery but also the enzymes necessary for target cleavage^8^, and in some cases for the processing of crRNAs^49^.

Our work revealed several negative associations between Cas subtypes. These may have selective or neutral causes. Some systems may be functionally very similar. In this case, the presence of the two systems in the genome is redundant and one of them is expected to be lost by drift. Some genomes have several clusters of the same Cas subtype (134 out of 2286 genomes with *cas* clusters), of which most (92) are type I and are usually in complex loci. Systems of the same subtype among Type II systems rarely co-occur (3 cases) which shows that these systems may be compatible, and their low co-occurrence might be explained by loss by drift. Alternatively, two systems may not work well together, e.g. because they compete for a substrate or because their mechanisms interfere in a deleterious manner. In this case, if a system is acquired by a genome and is incompatible with another already functioning in the cell, one expects selection for the deletion of one of them. This should result in rapid deletion of *cas* clusters, because this prevents CRISPRs from functioning. Such a mechanism may result in a CRISPR distant from functional *cas* clusters (when they are of a different type). Such negative interactions could be behind the peculiar system preventing the horizontal transfer of type I-F system in *E. coli* that encode the type I-E system ^50^. In this case, CRISPRs of type I-F contain spacers matching sequences of the cognate (absent) type I-F cas genes^50^. Upon acquisition of such system, spacers guide the incoming I-F CRISPR-Cas system to degrade incoming DNA and thus preventing acquisition of such systems^51^. The existence of such a mechanism suggests selection for preventing simultaneous presence of these two systems in the same genome.

The influx of novel CRISPR-Cas systems in genomes by mobile elements leads to co-existence of different systems in the same genome. Since transfer tends to accumulate in a small number of regions of the chromosome^52^, this leads to the accumulation of certain defence systems in the chromosome^53,54^. When such associations increase the immune competence of the cells, they may evolve to become integrated functional systems. This may explain why higher than expected co-occurrences between certain Cas subtypes were more frequent than the inverse, and why the multiple systems are often contiguous in single complex CRISPR-Cas loci. These complex organizations can reflect functional associations between CRISPR-Cas systems of various types, as suggested by the presence of a single adaptation module in many systems, and similar CRISPR repeats across the locus. Type III-B and type I-F are positively associated in Proteobacteria, and this could be explained by experimentally demonstrated ability of type III-B to process and use guide RNAs expressed from a type I-F CRISPR array^21^. The immunity provided by type III systems involves the production of an intracellular signal which activates a non-specific RNAse, Csm6. This mechanism can lead to cell death or dormancy when high levels of target mRNA are detected or when the target is mutated. As such, type III systems have been proposed to form a second line of defence able to block phage infection when type I systems fail^21^. It should be noted that this experimentally verified interaction between systems is based on two *cas* clusters that are not in a single locus. Hence, interactions between systems may start before they merge in a single complex locus. Together, these results suggests that associations between type I and type III systems combine the adaptation and interference functions of the former with a diversity of mechanisms associated with interference and abortive infective mechanisms of the latter. This suggests that the integration of multiple CRISPR on complex loci including multiple Cas systems can improve the immune defence of Prokaryotes against infection by mobile elements.

Supplementary Data are available at NAR online.

