## Supplementary Materials for "Co-occurrence of multiple CRISPRs and *cas* clusters suggests epistatic interactions"

#### Supplementary text 1

**Typing CRISPR arrays from metagenomes.**
CRISPRs can be detected in metagenomes[1–3]. It is challenging to associate Cas systems to them because the short size of most contigs in these datasets implicates that most arrays are split and in contigs apart from Cas clusters. We wanted to assess if our method could be used to assign subtypes to CRISPRs from metagenomes We downloaded 15 metagenomes from the HMP project[4] from 5 individuals for 3 body sites. We detected 806 spacers. We were able to assign a subtype to 92% of the spacers detected in the metagenomes. This number did not vary much between body sites from where samples were extracted with 83% of spacers typed in feces, 91% for buccal mucosa and 94% from gingiva. This shows that our method can be used to assign a subtype to spacers in metagenomes only using the array repeat (Supplementary Figure 5.C ).

**Methods for the metagenomes analysis.**
15 metagenomes were downloaded from the 5 individuals from HMP project (https://portal.hmpdacc.org/) with samples in three body sites: gingiva, feces, buccal mucosa (Supplementary Table 5). CRISPRs were detected using CRT with same parameters as the ones used in this study (default parameters except for –maxRL which was set to 50). We assigned subtypes by methods described for orphan arrays. We quantified the sequence similarity with all repeats of a database of repeats known to be associated with a specific Cas system using a global alignment with no gap end penalty and equal gap creation and extension penalties (-3) and took the best hit among those scores. If the identity score was higher than 72%, we assigned to the repeat the subtype of the best hit.

#### Supplementary tables

**Supplementary Table 1:** Cas clusters with associated CRISPRs
Large table, in annex.

**Supplementary Table 2:** CRISPRs with associated Cas clusters
Large table, in annex.

**Supplementary Table 3:** Key results using CRISPR arrays with a minimum of 5 repeats

| Analysis (main text) | Analysis (only CRISPR with 5 or more repeats) |
|---|---|
| Most *cas* clusters are associated with CRISPRs, but many CRISPRs are not next to *cas* clusters | This trend remains unchanged: CRISPRs are in CRISPR-Cas loci (65%), distant (24%) or orphan CRISPRs (16%). |
| Orphan arrays are smaller than distant arrays, which are smaller than arrays in CRISPR-Cas loci | These trends remain significantly different when considering arrays with more than 5 repeats (Tukey HSD, all pairs, P<0.001). |

**Supplementary table 4:** Characterization of repeats associated to specific subtypes (CRISPR-Cas loci with a single *cas* cluster).

| Subtype | Number of repeats (mean) | Number of repeats (median) | Maximum number of repeats | Number of Cas genes (mean) | Number of systems | Frequency of Cas1 presence | Frequency of Cas2 presence |
|---|---|---|---|---|---|---|---|
| IA | 50.69 | 31.50 | 400 | 8.13 | 116 | 0.91 | 0.91 |
| IB | 43.63 | 28.00 | 482 | 7.14 | 409 | 0.89 | 0.89 |
| IC | 32.56 | 21.00 | 319 | 6.51 | 464 | 0.88 | 0.88 |
| ID | 42.76 | 26.50 | 134 | 6.31 | 62 | 0.73 | 0.73 |
| IE | 21.20 | 14.00 | 240 | 7.55 | 1262 | 0.94 | 0.94 |
| IF | 26.01 | 17.00 | 249 | 5.73 | 341 | 0.94 | 0.94 |
| IIA | 16.34 | 13.00 | 71 | 4.00 | 180 | 0.99 | 0.99 |
| IIB | 27.21 | 28.00 | 46 | 3.79 | 14 | 0.93 | 0.93 |
| IIC | 23.33 | 19.00 | 113 | 3.02 | 206 | 1.00 | 1.00 |
| IIIA | 20.59 | 17.00 | 277 | 7.38 | 250 | 0.67 | 0.67 |
| IIIB | 23.11 | 14.00 | 237 | 6.29 | 168 | 0.50 | 0.50 |
| IIIC | 28.35 | 21.00 | 81 | 6.46 | 26 | 0.23 | 0.23 |
| IIID | 20.44 | 10.00 | 184 | 6.71 | 114 | 0.46 | 0.46 |
| IU | 36.38 | 20.00 | 589 | 4.73 | 81 | 0.70 | 0.70 |
| IV | 9.67 | 8.00 | 16 | 4.67 | 3 | 0.00 | 0.00 |
| V | 16.47 | 10.00 | 55 | 3.67 | 15 | 1.00 | 1.00 |
| VI | 10.89 | 10.50 | 39 | 1.00 | 36 | 0.00 | 0.00 |

**Supplementary table 5:** Metagenomes used in the study

| sample_id | subject_id | sample_body_site | subject_gender | file_id |
|---|---|---|---|---|
| 3674d95cd0d27e1de94ddf4d2efbe372 | 6788b769d0b62444378462474da09823 | gingiva | female | 91319e642fdd8a6e3b059cfb05bd6b68 |
| 3674d95cd0d27e1de94ddf4d2e6c41e0 | 6788b769d0b62444378462474da09823 | buccal mucosa | female | 91319e642fdd8a6e3b059cfb05c56868 |
| 3674d95cd0d27e1de94ddf4d2eeadee3 | 6788b769d0b62444378462474da09823 | feces | female | 91319e642fdd8a6e3b059cfb05bd5d02 |
| 3674d95cd0d27e1de94ddf4d2e06baea | 6788b769d0b62444378462474da09a28 | gingiva | male | 91319e642fdd8a6e3b059cfb05b11d56 |
| 3674d95cd0d27e1de94ddf4d2eeb0178 | 6788b769d0b62444378462474da09a28 | feces | male | 91319e642fdd8a6e3b059cfb05b7b659 |
| 3674d95cd0d27e1de94ddf4d2ec7ae18 | 6788b769d0b62444378462474da09a28 | buccal mucosa | male | 91319e642fdd8a6e3b059cfb05c2a4f3 |
| e2559e04fcd73935a7d7b917905184c2 | 6788b769d0b62444378462474da0a6c5 | gingiva | male | 91319e642fdd8a6e3b059cfb05b47d8c |
| e2559e04fcd73935a7d7b91790529c9d | 6788b769d0b62444378462474da0a6c5 | buccal mucosa | male | 91319e642fdd8a6e3b059cfb05c56568 |
| 3674d95cd0d27e1de94ddf4d2ec6161e | 6788b769d0b62444378462474da0a6c5 | feces | male | 91319e642fdd8a6e3b059cfb05b6dcab |
| 3674d95cd0d27e1de94ddf4d2ec6d386 | 6788b769d0b62444378462474da0b8a4 | feces | female | 91319e642fdd8a6e3b059cfb05ba5f2b |
| e2559e04fcd73935a7d7b917902c93d2 | 6788b769d0b62444378462474da0b8a4 | gingiva | female | 91319e642fdd8a6e3b059cfb05d32a9c |
| 3674d95cd0d27e1de94ddf4d2eb7bfc8 | 6788b769d0b62444378462474da0b8a4 | buccal mucosa | female | 91319e642fdd8a6e3b059cfb05e99f1e |
| 3674d95cd0d27e1de94ddf4d2e60bb7c | 6788b769d0b62444378462474da0c49e | feces | female | 91319e642fdd8a6e3b059cfb05c32d40 |
| 3674d95cd0d27e1de94ddf4d2efcbc50 | 6788b769d0b62444378462474da0c49e | gingiva | female | 91319e642fdd8a6e3b059cfb05b1df73 |
| e2559e04fcd73935a7d7b917901cbc39 | 6788b769d0b62444378462474da0c49e | buccal mucosa | female | 91319e642fdd8a6e3b059cfb05cb6467 |

### Supplementary Figures

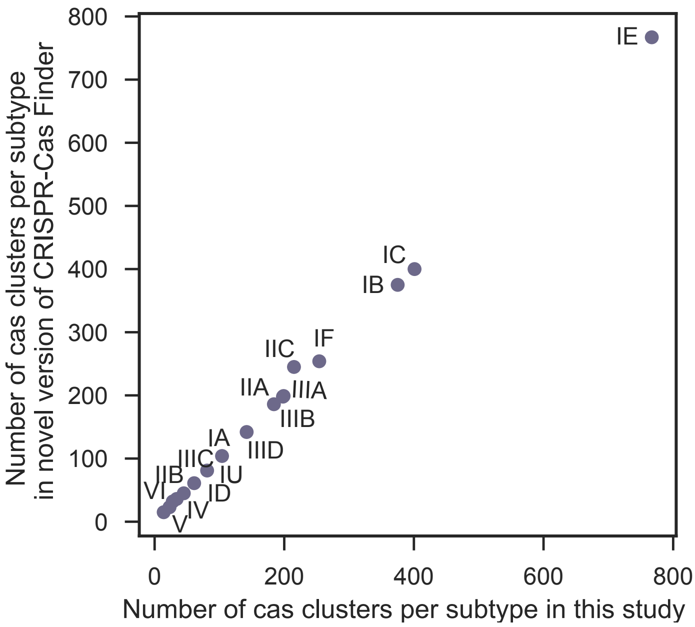


**Supplementary Figure 1: Comparison of the number of *cas* clusters identified with versions 0.9 and 1.0 of CRISPR-Cas Finder[5].**

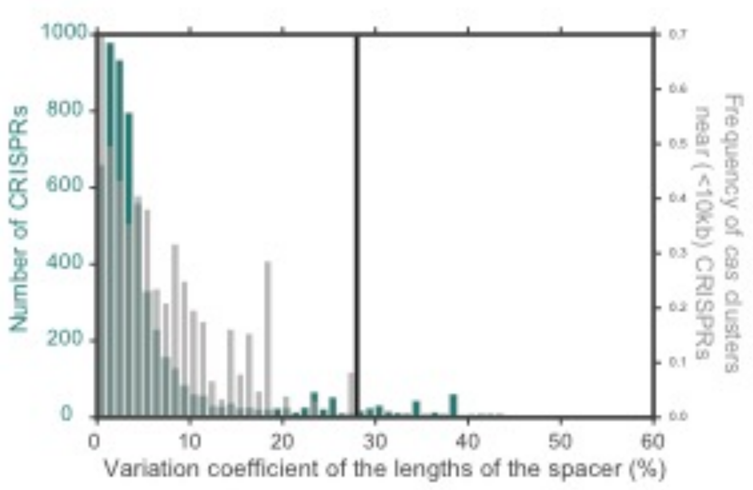


**Supplementary Figure 2: Detection of CRISPR-arrays: removing false elements.**
In green, the distribution of the coefficient of variation of the size of spacers of CRISPR arrays, in grey the observed frequencies of CRISPR with *cas* clusters within 10kb. The chosen coefficient of variation (in %) threshold of 28 is marked by a vertical line.

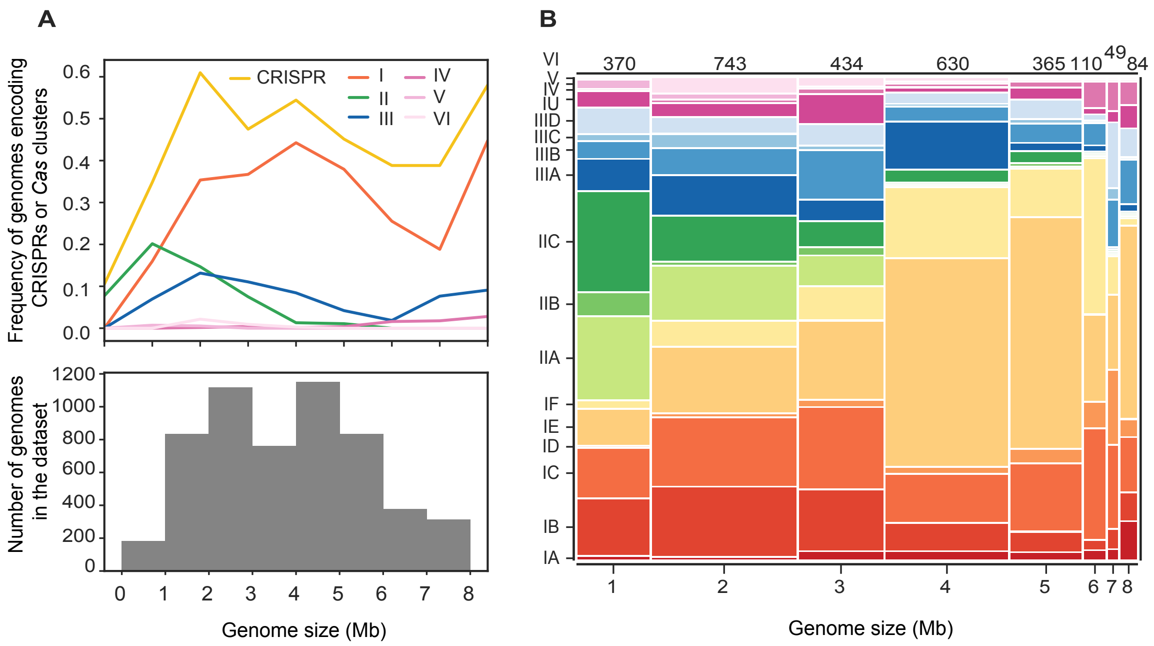


**Supplementary Figure 3 : Frequency of CRISPR arrays, Cas clusters and subtypes in function of genome size.**

**A.** Top panel represents the frequency of genomes carrying a system within the genome size range. Bottom panel represents the distribution of genomes according to genome size range. **B.** Frequency of subtypes in function of genome size range**.** Each color represent a subtype. X axis represents the genome size in Mb.

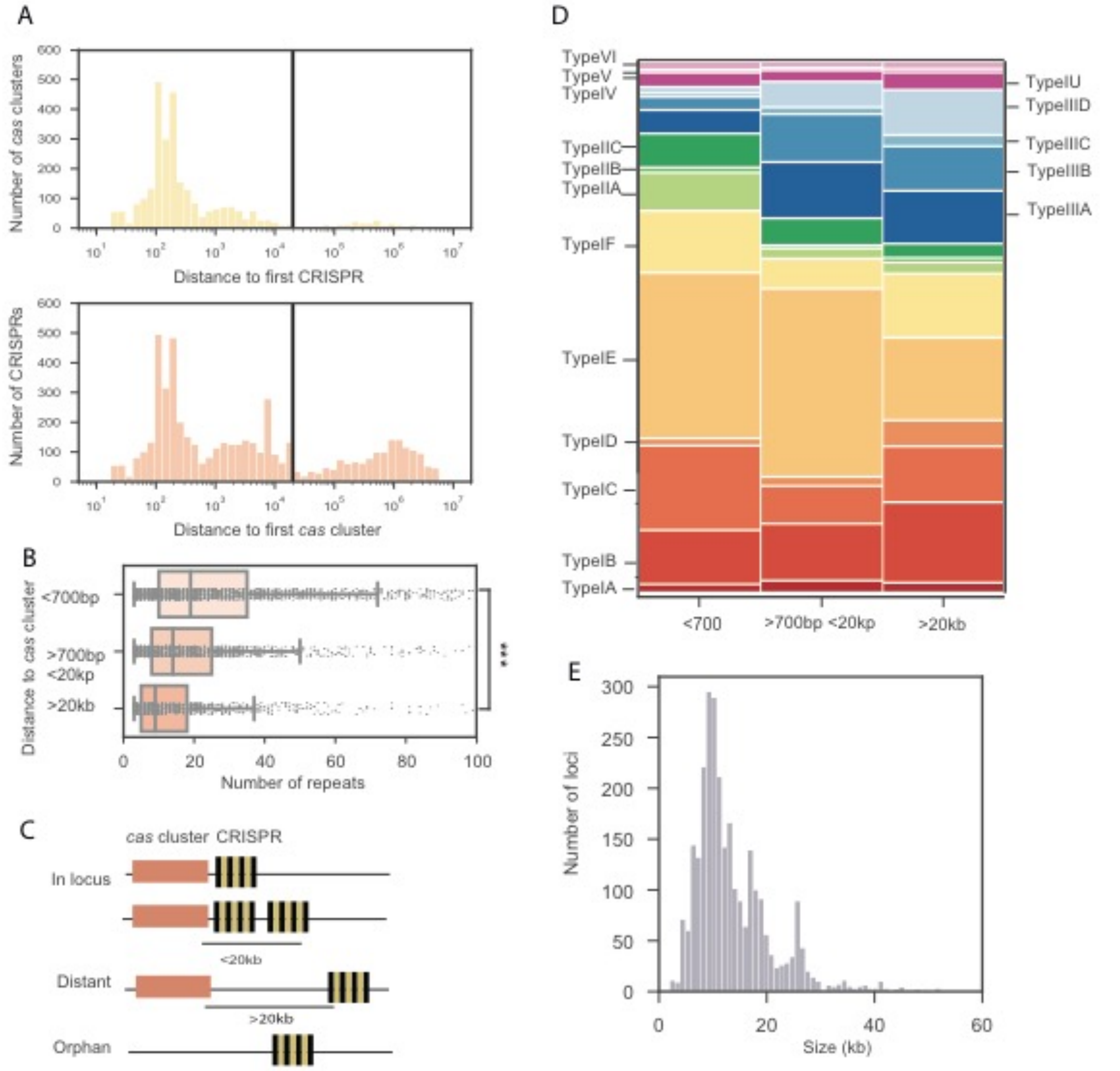


**Supplementary Figure 4: Comparison of three groups of CRISPR arrays and Cas clusters**

**A.** Distance between the CRISPR and the closest *cas* cluster (for CRISPR) and vice versa (for *cas* cluster). Vertical line corresponds to the distance cut-off (20kb). **B**. Boxplot of the number of repeats in different groups of CRISPR-arrays (ANOVA, P<0.001). **C.** Schematics of association of Cas clusters and CRISPR arrays **D.** Subtypes associated to the closest Cas clusters in the different groups. **E.** Histogram of the size of CRISPR-Cas loci.

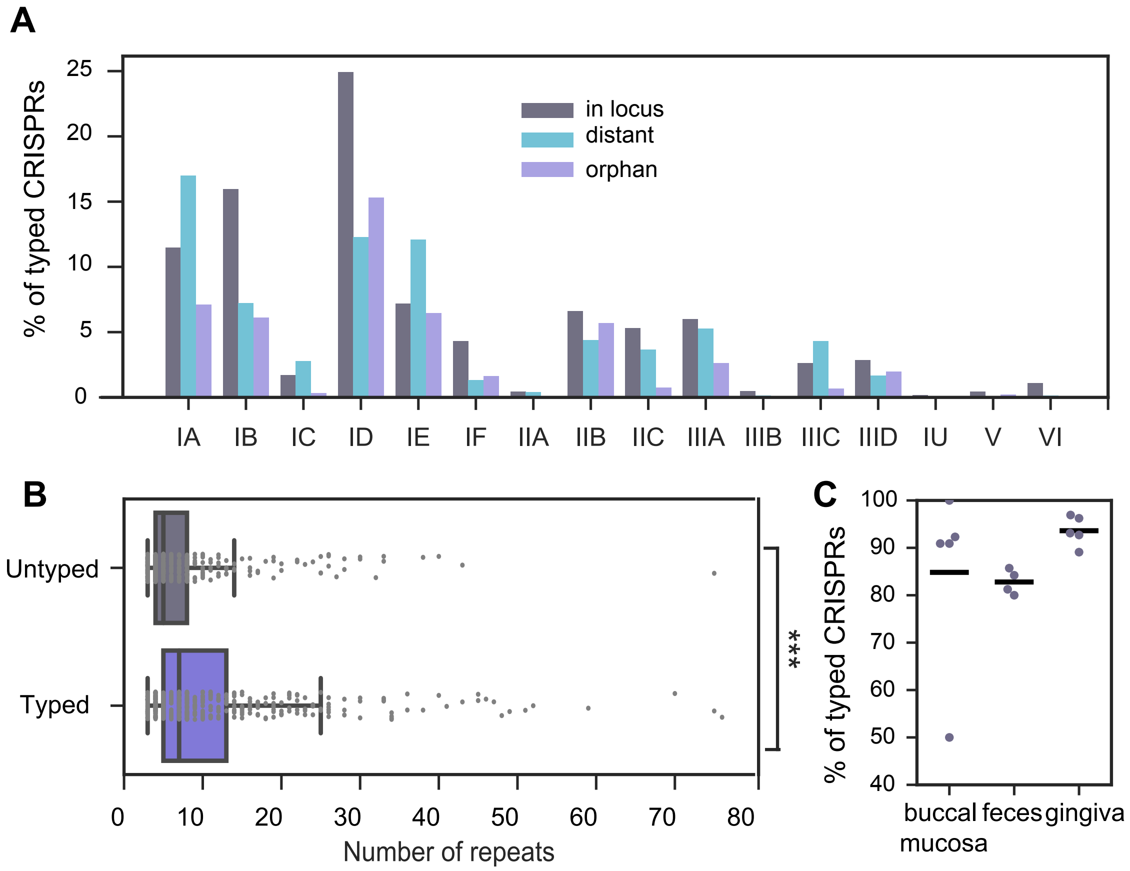


**Supplementary Figure 5 : Typing CRISPR arrays using best hit methods.**

**A.** Subtypes abundance in CRISPR arrays. CRISPR subtype was determined by the subtype of the associated Cas cluster for the "in loci". Distant and orphan CRISPR arrays were subtyped using the method described in the article, only based on their repeats. **B**. Boxplot of the number of repeats within untyped and typed orphan CRISPR arrays. Mann Whitney, P<0.001). X axis was cut at 100 for better visualization. **C**. Percentage of typed arrays of metagenomes

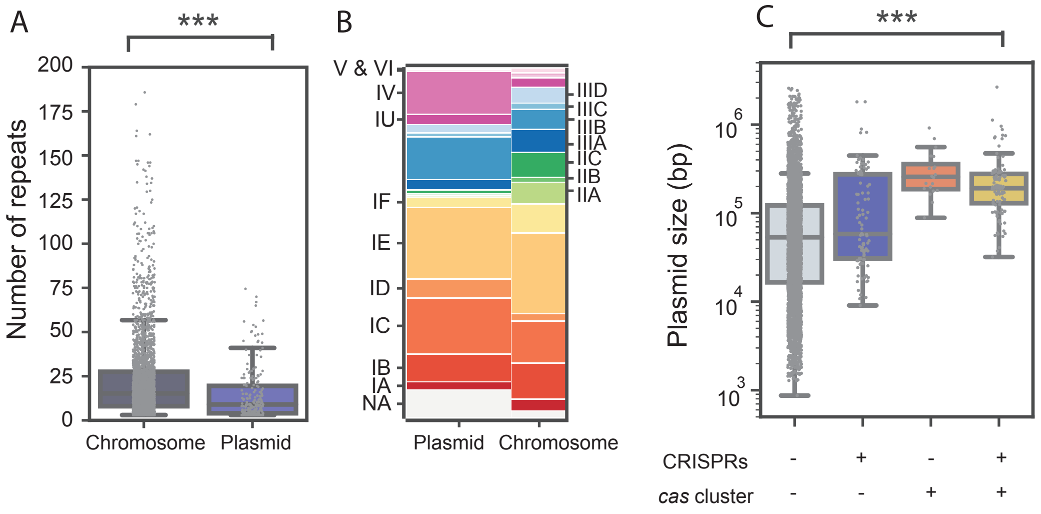


**Supplementary Figure 6 : Plasmids and CRISPR-Cas systems**

**A.** Boxplot of the number of repeats within chromosomal and plasmidic CRISPRs (Mann Whitney, P<0.001) **B.** Cas subtypes of systems identified in plasmids. Frequency of chromosomal and plasmid Cas subtypes (test on the independence of variables in a contingency table $\chi^2$, P<0.001). NA means that no specific subtype could be assigned. **C.** Boxplot of plasmid size in function of the presence of CRISPR-Cas systems (ANOVA, P<0.001).

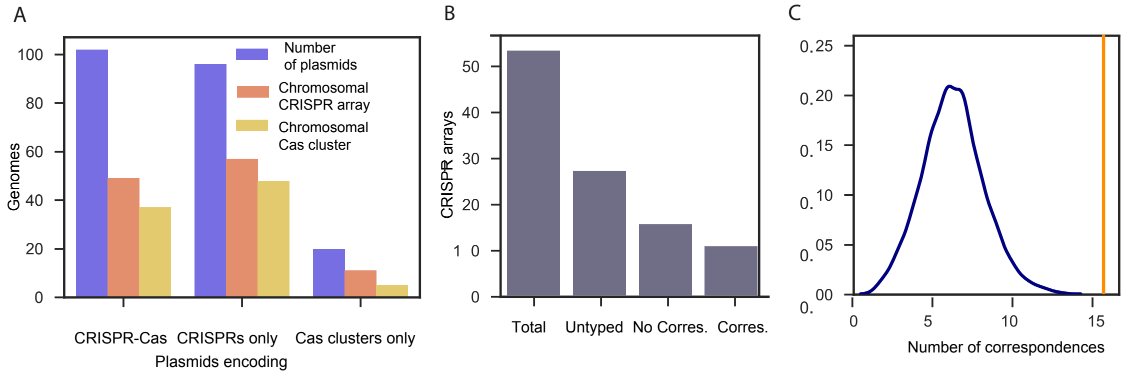


**Supplementary Figure 7: Subtype associations between plasmidic CRISPR arrays and associated chromosomal Cas clusters.**

**A. Number of plasmids with CRISPR-Cas (left), CRISPRs-only (center) of Cas-only (right).** The blue bar indicates the number of plasmids. The orange bar indicates the number of these plasmids that are in genomes whose chromosomes have a CRISPR and the bar in yellow those in genomes where the chromosome encodes a Cas system. **B.** Subtype correspondence between plasmidic CRISPR array and chromosomal Cas clusters. Correspondence was defined as such: If the assigned subtype of the array matched at least one of the Cas clusters present in the chromosome **C**. Control for plasmidic CRISPR array subtype correspondence to Cas cluster chromosomal one. Vector corresponding to the CRISPR subtypes was randomized and number of correspondence with the associated vector of Cas clusters was computed. The distribution of the number of correspondences for 1000 randomizations is presented. The number of observed correspondence in our dataset is represented by the vertical orange line.
